# Narcolepsy risk loci are enriched in immune cells and suggest autoimmune modulation of the T cell receptor repertoire

**DOI:** 10.1101/373555

**Authors:** Hanna M Ollila, Eilon Sharon, Ling Lin, Nasa Sinnott-Armstrong, Aditya Ambati, Ryan P Hillary, Otto Jolanki, Juliette Faraco, Mali Einen, Guo Luo, Jing Zhang, Fang Han, Han Yan, Xiao Song Dong, Jing Li, Jun Zhang, Seung-Chul Hong, Tae Won Kim, Yves Dauvilliers, Lucie Barateau, Gert Jan Lammers, Rolf Fronczek, Geert Mayer, Joan Santamaria, Isabelle Arnulf, Stine Knudsen, May Kristin Lyamouri Bredahl, Per Medbøe Thorsby, Giuseppe Plazzi, Fabio Pizza, Monica Moresco, Catherine Crowe, Stephen K Van den Eeden, Michel Lecendreux, Patrice Bourgin, Takashi Kanbayashi, Rosa Peraita-Adrados, Francisco J Martínez-Orozco, Antonio Benetó, Jacques Montplaisir, Alex Desautels, Yu-Shu Huang, Poul Jennum, Sona Nevsimalova, David Kemlink, Alex Iranzo, Sebastian Overeem, Aleksandra Wierzbicka, Peter Geisler, Karel Sonka, Makoto Honda, Birgit Högl, Ambra Stefani, Fernando Morgadinho Coelho, Vilma Mantovani, Eva Feketeova, Mia Wadelius, Niclas Eriksson, Hans Smedje, Pär Hallberg, Per Egil Hesla, David Rye, Zerrin Pelin, Luigi Ferini-Strambi, Claudio L Bassetti, Johannes Mathis, Ramin Khatami, Adi Aran, Sheela Nampoothiri, Tomas Olsson, Ingrid Kockum, Markku Partinen, Markus Perola, Birgitte R Kornum, Sina Rueger, Juliane Winkelmann, Taku Miyagawa, Hiromi Toyoda, Seik Soon Khor, Mihoko Shimada, Katsushi Tokunaga, Manuel Rivas, Jonathan K Pritchard, Neil Risch, Zoltan Kutalik, Ruth O’Hara, Joachim Hallmayer, Chun Jimmie Ye, Emmanuel Mignot

## Abstract

Type 1 narcolepsy (T1N) is a neurological condition, in which the death of hypocretin-producing neurons in the lateral hypothalamus leads to excessive daytime sleepiness and symptoms of abnormal Rapid Eye Movement (REM) sleep. Known triggers for narcolepsy are influenza-A infection and associated immunization during the 2009 H1N1 influenza pandemic. Here, we genotyped all remaining consented narcolepsy cases worldwide and assembled this with the existing genotyped individuals. We used this multi-ethnic sample in genome wide association study (GWAS) to dissect disease mechanisms and interactions with environmental triggers (5,339 cases and 20,518 controls). Overall, we found significant associations with HLA (2 GWA significant subloci) and 11 other loci. Six of these other loci have been previously reported (*TRA*, *TRB*, *CTSH*, *IFNAR1*, *ZNF365* and *P2RY11*) and five are new (*PRF1*, *CD207*, *SIRPG*, *IL27* and *ZFAND2A)*. Strikingly, in vaccination-related cases GWA significant effects were found in *HLA*, *TRA,* and in a novel variant near *SIRPB1*. Furthermore, *IFNAR1* associated polymorphisms regulated dendritic cell response to influenza-A infection in vitro (p-value =1.92*10^−25^). A partitioned heritability analysis indicated specific enrichment of functional elements active in cytotoxic and helper T cells. Furthermore, functional analysis showed the genetic variants in *TRA* and *TRB* loci act as remarkable strong chain usage QTLs for *TRAJ*24 (*p-value = 0.0017), *TRAJ*28* (p-value = 1.36*10^−10^) and *TRBV*4-2* (p-value = 3.71*10^−117^). This was further validated in TCR sequencing of 60 narcolepsy cases and 60 DQB1*06:02 positive controls, where chain usage effects were further accentuated. Together these findings show that the autoimmune component in narcolepsy is defined by antigen presentation, mediated through specific T cell receptor chains, and modulated by influenza-A as a critical trigger.

## Main Text

Type 1 narcolepsy (T1N) is a sleep disorder that affects 1/3,000 individuals across ethnic groups^1-3^. Onset is typically in childhood through early adulthood. Symptoms are caused by the destruction of hypocretin/orexin neurons, a small neuronal subpopulation of the hypothalamus^4^. Although the disease is considered autoimmune, the exact mechanism leading to hypocretin cell death is still unclear. Indeed, T1N is strongly associated with alleles encoding the heterodimer DQ0602 haplotype (HLA-*DQA1*01:02*~*DQB1*06:02*, 97% vs. 25%) across ethnic groups^5,6^. Other loci previously associated with the disease include T cell receptor (TCR) loci alpha (*TRA*) and beta (*TRB*), receptors of HLA-peptide presentations, and other autoimmune associated genes (*CTSH, P2RY11, ZNF365, IFNAR1* and *TNFSF4*)^7-10^.

Triggers of T1N point to the immune system, including influenza and Streptococcus Pyogenes infections^9,11,12^, as well as immunization with Pandemrix^®^, an influenza-A vaccine developed specifically against the H1N1 “swine flu” strain 13-20 suggest a strong environmental modifier of disease risk for narcolepsy. Increased T1N incidence following the Pandemrix^®^ vaccination was first seen in Northern Europe^13-20^ with 8-fold increase in incidence in (0.79/100,000 to 6.3/100,000) in children. The specificity was striking, as increased T1N was later detected in all countries where Pandemrix^®^ was used, whereas countries using other pH1N1 vaccine brands did not detect vaccination-associated increases in incidence^13-22^.

Despite the genetic and epidemiological evidence for T1N being an immune-system mediated disease, only a few genetic risk factors have been found or characterized so far. Furthermore, the functional consequence of these variants has remained unstudied. Therefore, we examine and characterize genetic factors for T1N across multiple ethnic groups in a sample three times larger than earlier studies finding novel mechanisms how these variants affect RNA expression and T cell receptor chain usage. Our novel findings show that the autoimmune component in narcolepsy is defined by antigen presentation, mediated through specific T cell receptor chains, and modulated by influenza-A as a critical trigger.

## Results

### GWAS discovers five novel risk loci for narcolepsy

To discover novel narcolepsy loci, we first meta-analyzed a large multiethnic cohort of 5,339 T1N cases and 20,518 controls consisting of samples from nine independent cohorts across three ethnic groups. In addition to the strongest associations in the HLA locus (minimum p-value<10^−216^), we discovered additional 228 genome-wide significant SNPs with no evidence of genomic inflation^23^ (λ=1.06) (meta-analysis p-value < 5×10^−8^; **Fig. 1)**. These results confirmed six out of eight previously identified loci (*TRA*, *TRB, CTSH, IFNAR1, ZNF365* and *P2RY11*), and identified five novel loci near *CD207, SIRPG, IL27, ZFAND2A* and *PRF1* (**Fig. 1**, **Table 1, Supplementary Figs. 1-2**). Further fine-mapping suggested more than one signal in *TRB, ZNF365, TRA, SIRPG* and *IFNAR1* loci (Supplementary information). Furthermore, a GCTA gene based test^24^ showed association with three known autoimmune or inflammatory disease genes with *GPR25^25,26^*, *C1ORF106* ^27^ and *PD-1*28,29, suggesting that additional variants remain to be discovered using larger sample sizes (see **Supplementary Tables 1-3**) doubling the number of variants in T1N.

**Fig. 1.**
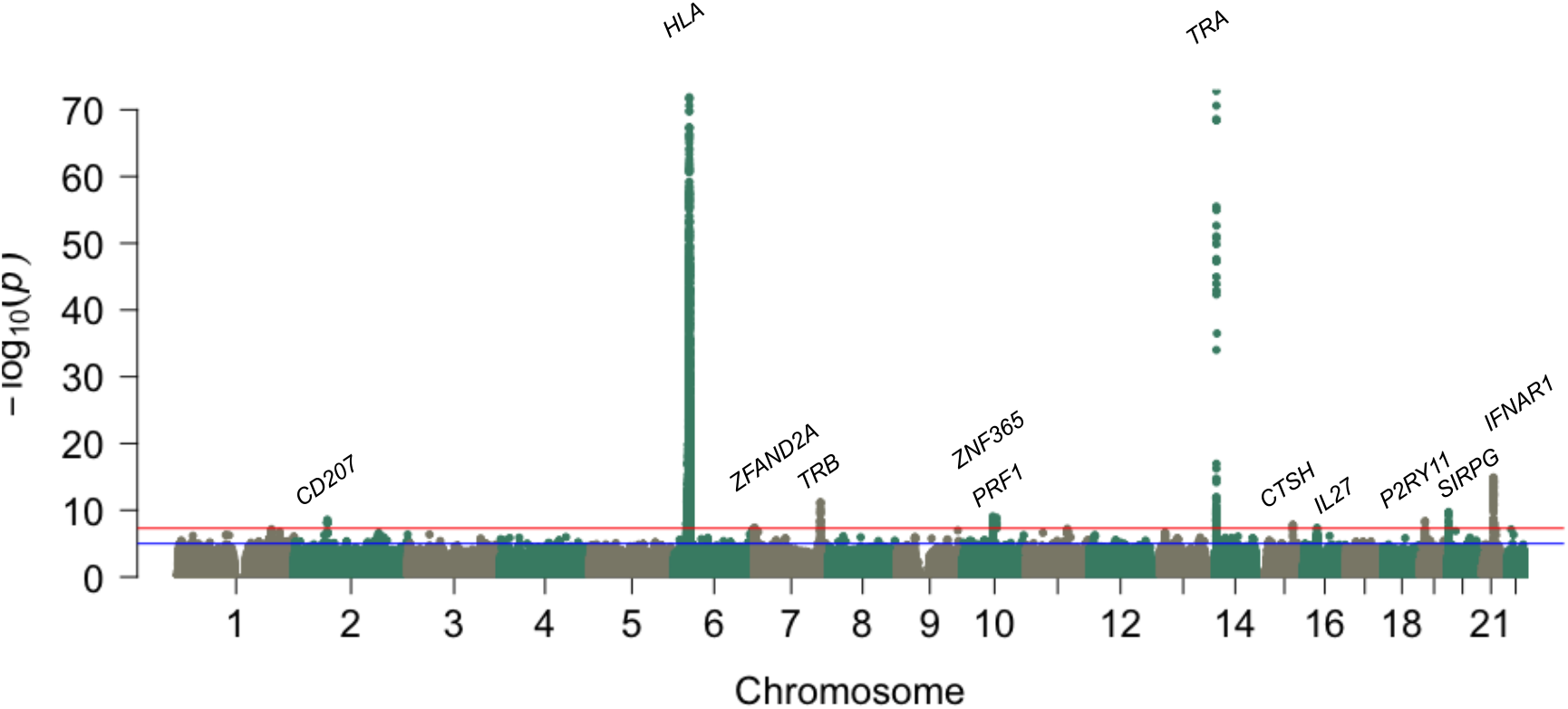
Multi ethnic genetic analysis of type 1 narcolepsy. Multi-ethnic analysis conducted in 5,339 cases and 20,518 controls reveals genome-wide significant associations in 11 loci plus HLA. The x-axis shows genomic location by chromosome and the y-axis shows −log_10_ p-values. Red horizontal line indicates genome-wide significant p-value threshold of 5*10^−8^. P-values smaller than 10^−75^ were set to 10^−75^ (HLA locus has many SNPs with p-value<10^−216^).

**Table 1.** Genome-wide significant associations observed in T1N across ethnic groups.

| Closest Gene | chr | rsid | pos | non-coded allele | coded allele | p-value | coded af | OR [CI lower - upper] | beta | se |
| --- | --- | --- | --- | --- | --- | --- | --- | --- | --- | --- |
| CD207 (Langerin) | 2 | rs13383830 | 71058306 | T | C | 2.65E-09 | 0.078 | 1.285 [1.184-1.396] | 0.251 | 0.042 |
| ZFAND2 | 7 | rs75674288 | 1195322 | A | C | 4.05E-08 | 0.913 | 0.778 [0.711-0.851] | -0.251 | 0.046 |
| TRB | 7 | rs1008599 | 142038782 | A | G | 6.63E-12 | 0.332 | 0.813 [0.767-0.862] | -0.207 | 0.03 |
| ZNF365 | 10 | rs4237304 | 64407845 | C | T | 8.40E-10 | 0.824 | 1.233 [1.154-1.319] | 0.21 | 0.034 |
| PRF1 | 10 | rs35947132 | 72360387 | G | A | 1.40E-09 | 0.04 | 0.570 [0.475-0.684] | -0.562 | 0.093 |
| TRA | 14 | rs1154155 | 23002684 | T | G | 1.48E-73 | 0.255 | 1.643 [1.559-1.733] | 0.497 | 0.027 |
| CTSH | 15 | rs34593439 | 79234957 | G | A | 1.44E-08 | 0.09 | 1.246 [1.154-1.345] | 0.22 | 0.039 |
| IL27 | 16 | rs200840505 | 28539396 | GTGTGTA | G | 4.70E-08 | 0.281 | 0.849 [0.801-0.901] | -0.163 | 0.03 |
| P2YR11 | 19 | rs34849604 | 10229098 | T | TG | 4.26E-09 | 0.537 | 1.232 [1.148-1.323] | 0.209 | 0.036 |
| SIRPG | 20 | rs6110697 | 1615661 | T | C | 1.83E-10 | 0.74 | 1.206 [1.138-1.28] | 0.188 | 0.03 |
| IFNAR1 | 21 | rs2409487 | 34684958 | C | T | 1.23E-15 | 0.754 | 1.214 [1.158-1.273] | 0.194 | 0.024 |

Leading SNP of loci associated with T1N at a genome wide significant level ( $p\text{-value} < 5 \times 10^{-8}$ ). Heterogeneity p-value is calculated between the nine cohorts in this study. Altogether 228 variants were significantly associated with T1N. Associations tested using SNPtest, and META with fixed effects test statistics are shown<sup>99,107</sup>. Positions are shown for genome build human genome build 37 (GRCh37/hg19).

Next we examined the genetic architecture of T1N by calculating the narrow sense heritability explained by the typed variants. GCTA estimated the observed scale heritability to be h2_SNP_[ci]=0.403 [0.015] ^30^ and the population heritability to be h2_SNP_[ci]= 0.231 [0.0088] assuming a prevalence estimate of 0.03%^1,2^. One third of observed heritability was mediated by genetic variation within the extended MHC region and similar to other pediatric autoimmune diseases ^31^.

### Narcolepsy shares variants with autoimmune diseases

We next examined genome-wide shared genetic correlation with other traits excluding variants at the extended HLA locus. ^32^ Note that we performed this analysis using samples of Whites as reflecting the genetic makeup of the population for which public data is available. The strongest correlations were seen between T1N and autoimmune diseases (Wilcox signed rank p-value = 0.031). Of all autoimmune traits examined using LD Score Regression^33^, the shared heritability was largest with type-1 diabetes (T1D) (r_g_=0.3261 (se=0.1015), p-value = 0.0013).

We next examined whether genome wide significant T1N associations are shared with other autoimmune diseases, suggesting shared mechanisms at single loci. Significant associations in T1N were compared with autoimmune disease associations using published studies and GWAS central^37-39^. Most notably, co-localization of signals using coloc anaysis ^40^ was found at *IL27* between T1N and both ankylosing spondylitis (posterior probability [pp] = 0.96) and Crohn’s disease (pp=0.93).

We also discovered strong overlap between T1N and T1D at *CTSH* pp=0.998 and *SIRPG* pp=0.999, as well as evidence for partial sharing at *IL27* pp=0.71, while signals were independent for *P2RY11* (pp=0.02). T1D is also the only autoimmune trait besides narcolepsy where any association was seen near the TRA locus, although the T1D signal (rs7145202, beta = 0.1, p-value = 4*10^−6^)^41^ is independent from the narcolepsy signal (r^2^<0.5) and located ~100 kb upstream of the TRA loci per se. While previous studies have shown either a small increase or no increased risk for autoimmune diseases in T1N patients, ^34-36^ we found statistical evidence of global genetic correlation between T1N and other autoimmune diseases and co-localization of individual associations.

### Genetics of vaccination-triggered narcolepsy

We have previously shown that both influenza infections and, in rare cases, immunization with Pandemrix^®^ can trigger narcolepsy^13,18,19,42,43^. The baseline for narcolepsy in unvaccinated vs. Pandemrix^®^ vaccinated individuals was 0.7/100,000 vs. 9/100,000 person years with on average 10-fold increase in risk ^13,18,19,42-44^. We therefore recruited Pandemrix^®^ vaccination-related narcolepsy cases in five countries and examined the genetic load for narcolepsy (**Table 2**). All Pandemrix^®^ vaccination cases were carriers also for HLA-DQB1*06:02. Weighted genetic risk score (GRS) excluding HLA showed a strong association in Pandemrix^®^ vaccination related narcolepsy in each sub cohort (p<0.01 for all cohorts) and with combined vaccination related narcolepsy sample (p-value = 7.96*10^−10^). (Table 2, and Supplementary Table 8 and Supplementary Figs. 3-4).

**Table 2.** Locus specific (from Table 1) and Genome-Wide significant associations observed in vaccination-triggered T1N cases.

| Closest Gene | chr | rsid | pos | non-coded<br>allele | coded<br>allele | p-value | OR | beta | se | p-value<br>heterogeneity |
| --- | --- | --- | --- | --- | --- | --- | --- | --- | --- | --- |
| CD207<br>(Langerin) | 2 | rs13383830 | 71058306 | T | C | 0.757 | 1.106<br>[0.584 - 2.097] | 0.101 | 0.326 | 0.421 |
| ZFAND2 | 7 | rs75674288 | 1195322 | A | C | 4.92E-04 | 3.189 [1.66-6.122] | 1.160 | 0.333 | 0.673 |
| TRB | 7 | rs1008599 | 142038782 | A | G | 0.099 | 0.798<br>[0.61 - 1.043] | -0.226 | 0.137 | 0.822 |
| ZNF365 | 10 | rs4237304 | 64407845 | C | T | 0.410 | 1.116<br>[0.85 - 1.450] | 0.110 | 0.133 | 0.948 |
| PRF1 | 10 | rs35947132 | 72360387 | G | A | 4.48E-04 | 0 [0 - inf] | -10.879 | 50.123 | 1* |
| TRA | 14 | rs1154155 | 23002684 | T | G | 1.58E-13 | 2.531<br>[1.978 - 3.239] | 0.929 | 0.126 | 0.365 |
| CTSH | 15 | rs34593439 | 79234957 | G | A | 0.074 | 1.418<br>[0.967 - 2.079] | 0.349 | 0.195 | 0.614 |
| IL27 | 16 | rs200840505 | 28539396 | GTGTGTA | G | 0.318 | 0.834<br>[0.583 - 1.191] | -0.182 | 0.182 | 1** |
| P2YR11 | 19 | rs34849604 | 10229098 | T | TG | 0.024 | 1.515<br>[1.057 - 2.171] | 0.415 | 0.184 | 1** |
| SIRPG | 20 | rs6110697 | 1615661 | T | C | 0.050 | 1.336 [1.00-1.785] | 0.290 | 0.148 | 0.737 |
| IFNAR1 | 21 | rs2409487 | 34684958 | C | T | 0.720 | 1.077<br>[0.719 - 1.614] | 0.074 | 0.206 | 0.964 |
| SIRPB1-SIRPG | 20 | rs76958425 | 1602668 | C | T | 1.12E-08 | 2.491<br>[1.821 - 3.408] | 0.913 | 0.16 | 0.078 |

Association with vaccination related narcolepsy is shown for loci having genome wide significant association with T1N or those loci being
genome-wide significant with vaccination related narcolepsy. Associations tested using SNPtest or Chisq test (Irish) with meta-analysis
using META with fixed effects test statistics are shown<sup>99,107</sup>. Positions are shown for genome build human genome build 37

Similarly to GRS evidenced shared signal, we found GWA significant signal with HLA-DQB1*06:02, *TRA* rs1154155 and a variant between *SIRPB1*-*SIRPG* locus (rs76958425, OR=2.49 [1.82 - 3.41], p-value = 1.12*10-8, Table 2) not present in regular cases (rs76958425, p-value=0.15, beta = −0.0694, OR=0.93). The overall association of GRS and two shared loci indicate that vaccination related narcolepsy is fundamentally the same disorder as idiopathic T1N.

### Functional analyses highlight effects on immune cells

Analysis using GARFIELD^45^ showed the variants with p-value<0.00001 have a 5.9-fold enrichment for missense variants and 5.3 fold enrichment for 5’UTRs (**Fig.2, Supplementary Figs 5-7**). Further, many associated variants in Table 1 are in tight linkage with non-synonymous substitutions in the corresponding genes, such as variants in CTSH (rs2289702 G11R), TRA (rs1483979, F8L), PRF1 (rs35947132, A91V), SIRPG (rs6043409, V263A), CD207 (rs13383830, N288D and rs57302492, K313I, r2 =1) and IL27 (rs181206 L119P) as well as variants marking different HLA-alleles.

**Fig. 2.**
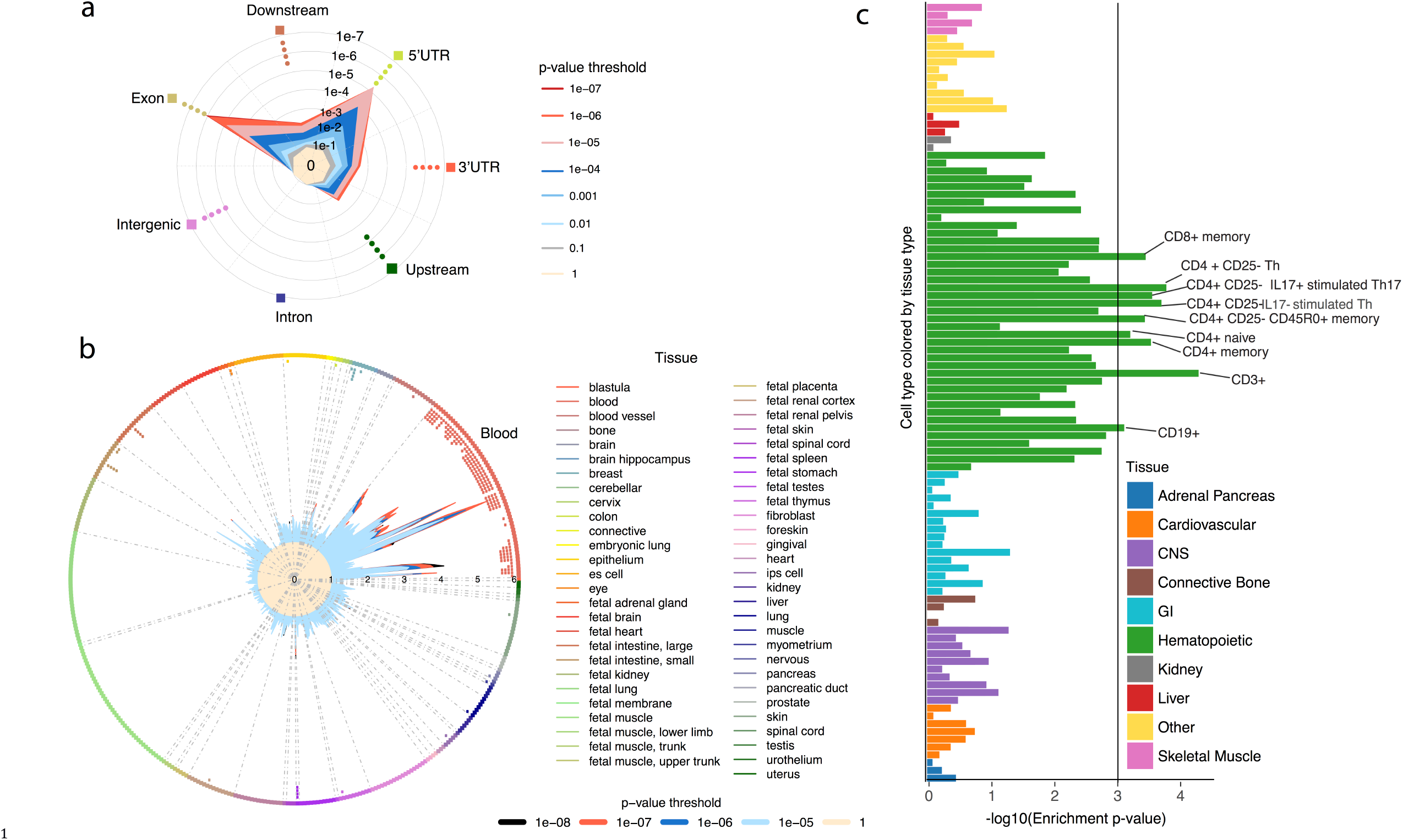
Narcolepsy risk variants are enriched in immune cells and for missense variants. a) GARFIELD analysis of narcolepsy associated variants shows a 6 fold enrichment for exon variants and a 5.2 fold enrichment in 5’UTRs located variants. b) overall enrichment in DNA hypersensitivity regions is seen specifically in circulating hematopoietic (blood) cells c) Epigenome roadmap data shows that the majority of narcolepsy heritability is enriched in hematopoietic cell lineages, with changes most pronounced in immune cells notably T helper and cytotoxic cells. Statistically significant enrichment is marked with a line corresponding to an Benjamin Hochberg enrichment p-value = 0.001.

We confirmed that variants within *CTSH* are also important in the predisposition of T1N. Among immune cells, *CTSH* is only expressed in Class II positive antigen presenting cells (B cells, dendritic cells and monocytes), and is known to process antigen for HLA presentation, thus furthering a role for HLA-DQ presentation in T1N. Of note, we also observed a sub threshold association with another cathepsin gene, *CTSC* (rs3888798, C allele frequency =0.06), OR = 1.276 [1.169-1.394] p-value =5.8*10^−8^), which was not associated with vaccination related narcolepsy (rs3888798, OR=0.76, p-value= 0.336).

In *PRF1*, the leading variant rs35947132 causes an amino acid change A91V that acts as a hypomorph and disrupts cytotoxicity of the immunological HLA class I synapse^46,47^. This relatively rare variant (allele frequency 0.03 in Whites) has been shown to prevent perforin, a protein expressed only by natural killer (NK) and cytotoxic (CD8^+^) T cells, to form functional complexes, thus preventing cytotoxic cells from destroying target cells^46,48^. These findings indicate direct involvement of cytotoxic T cells, most likely CD8^+^ T cells, in hypocretin cell destruction.

In addition, we discovered associations is in signal-regulatory protein gamma *SIRPG* (rs6110697, V263A) a receptor-type transmembrane glycoprotein known to interact with *CD47*, an anti-autophagy signal for the immune system that has shown success in cancer immunotherapy^59^. Although V263 is conserved in all SIRP family members, it is also located within an alternate exon. Unlike other members of the SIRP family, *SIRPG* is almost exclusively expressed in CD4^+^ and CD8^+^ T cells. Furthermore, the SNP is also a strong eQTL in thymus and whole blood^60^. Interestingly, vaccination-associated cases displayed an additional GWAS significant association with rs76958425, a strong QTL for *SIRPB1*, another SIRP family member known to interact with *CD47*. This association is not present in the overall narcolepsy sample (rs76958425, beta = - 0.0694138,OR=0.93, p=0.15). *SIRPB1* is mostly expressed in antigen presenting cells and has been shown to modulate neuronal killing in Alzheimer’s disease^61^, suggesting it could also be important for hypocretin-cell survival, though it may play a role in the modulation of T cell population survival.

One of the strongest novel factors associated with narcolepsy is rs2409487 in the *IFNAR1* gene, a gene mediating interferon α*/*β inhibition of virus replication type 1 interferon response associated with T1N. We observed that this SNP is a strong eQTL for *IFNAR1* expression in various tissues in GTEx^68^. In addition, a different lead variant (e.g. rs2284553) has been associated with other autoimmune diseases. IFNAR1 controls dendritic cell responses to viral infections, notably influenza A^69^. We therefore examined IFNAR1 expression in DC following H1N1 infection (PR8 delta NS1) finding that our predisposing SNP (rs2409487) is a major eQTL for this effect (p-value = 1.92*10^−25^, beta =0.140), and in perfect LD with the leading variant for the signal (rs6517159, D’=1, r^2^=0.995, coloc pp = 0.964 **Supplementary Fig. 8**). The findings suggest that rs2409487 in *INFAR1* mediates predisposition to T1N by modulating response to Influenza-A infection.

### Overlap of risk with cell-type specific chromatin regions

We examined whether associations with narcolepsy were enriched genome-wide on specific enhancer elements using stratified LD score regression on Epigenome Roadmap cell type specific annotations (n=216 cell types)^71^. Partitioned heritability by functional categories enriched in the hematopoietic cell lines (**Supplementary Fig. 2b and 2c, Supplementary Fig 8.**). Consistent with our model, association was driven by CD4^+^ T cells, with leading effects in CD3+ primary H3K27ac, CD4+/CD25-/IL17-PMA&ionomycin stimulated primary H3K4me1, and CD4+/CD25-primary H3K4me1 (each enriched over 35-fold in predicted heritability per SNP). Additional effects were seen in Th17 CD4^+^ T cells and CD8^+^ T cells, confirming the importance of these cell types in narcolepsy. Importantly, no enrichment was seen in neuronal cell types. While immune cells have been suggested to play a role in the predisposition to T1N^72^, these novel findings show that the effects are specific to both helper and cytotoxic T cells, and that individual variants genome-wide are substantially enriched in specific T cell lineages predisposing to T1N.

### Risk variants in T cell receptor loci modulate αβ T cell receptor repertoire

T1N is the only autoimmune disease with significant association in HLA and T cell receptor (TCR) loci (TRA and TRB). TCR molecules are formed through VDJ somatic recombination at the genomic level, a process that allows for substantial TCR sequence diversity. The recombinant T cell clones are later subjected to negative and positive selection in the thymus in order to optimize pathogen responses while avoiding auto-reactivity. As a consequence, most of TCR binding diversity is ensured by selection in the context of specific HLA molecules. TCRα and β chains heterodimerize to form biologically functional molecules that recognize peptides presented by the Major histocompatibility complex (MHC) encoded by the highly variable classical HLA genes. On one hand, T1N is associated with the DQB1*06:02 allele of the MHC class II β subunit and the highly linked DQA1*01:02 allele of MHC class II α subunit. On the other hand, T1N is strongly associated with TCR α and b chains. Notably, this association is also seen in cases with vaccination-triggered narcolepsy (**Table 2**). This suggests that T1N is directly linked with autoimmunity that is mediated by T-cell activation.

Clearly the strong association of T1N with the HLA locus will affect the presented epitope and the TCR repertoire^73^, however how does the association with TRA and TRB affect the TCR repertoire? In the TRB region, association peaks over 32 SNPs (from hg19 chr7:142025523-142248636) over a 22kb segment. The association signal within TRA locus spans over several J genes over 18 kb, with 5 SNPs (rs1154155, rs1483979, rs3764159, rs3764160) in perfect LD across ethnic groups. Among TRA SNPs, rs1483979, a SNP changing F8L in the peptide recognizing groove of CDR3 region of TRA J24 is an obvious candidate defining two J24 alleles we denote as J24*01 and J24*02 respectively. We next examined the effects of these SNPs on T-cell receptor V or J gene chain usage using RNA sequencing in 895 individuals^73^. Strikingly, rs1154155 with TRA J28 expression in total RNA sequencing from blood (p-value=1.36*10^−10^, beta = −0.212, **Fig. 3**) with the same lead variants that associated with narcolepsy and posterior probability for shared variant was pp=0.958 suggesting that rs1154155 in T1N predisposition mediates its action through effects on TRA J28 repertoire (See supplementary **Table S5** for all rs1154155 effects). J24 usage is also among the top associations for rs1154155 effects, although in this case correlation is opposite and the associated SNP increases usage (p-value=0.0017, beta=0.104, pp=0.54). Associations with multiple target variants within the same haplotype have been defined with complex traits with both regulatory and non-coding effects before and are likely to have a role in T1N predisposition^74^.

**Fig. 3.**
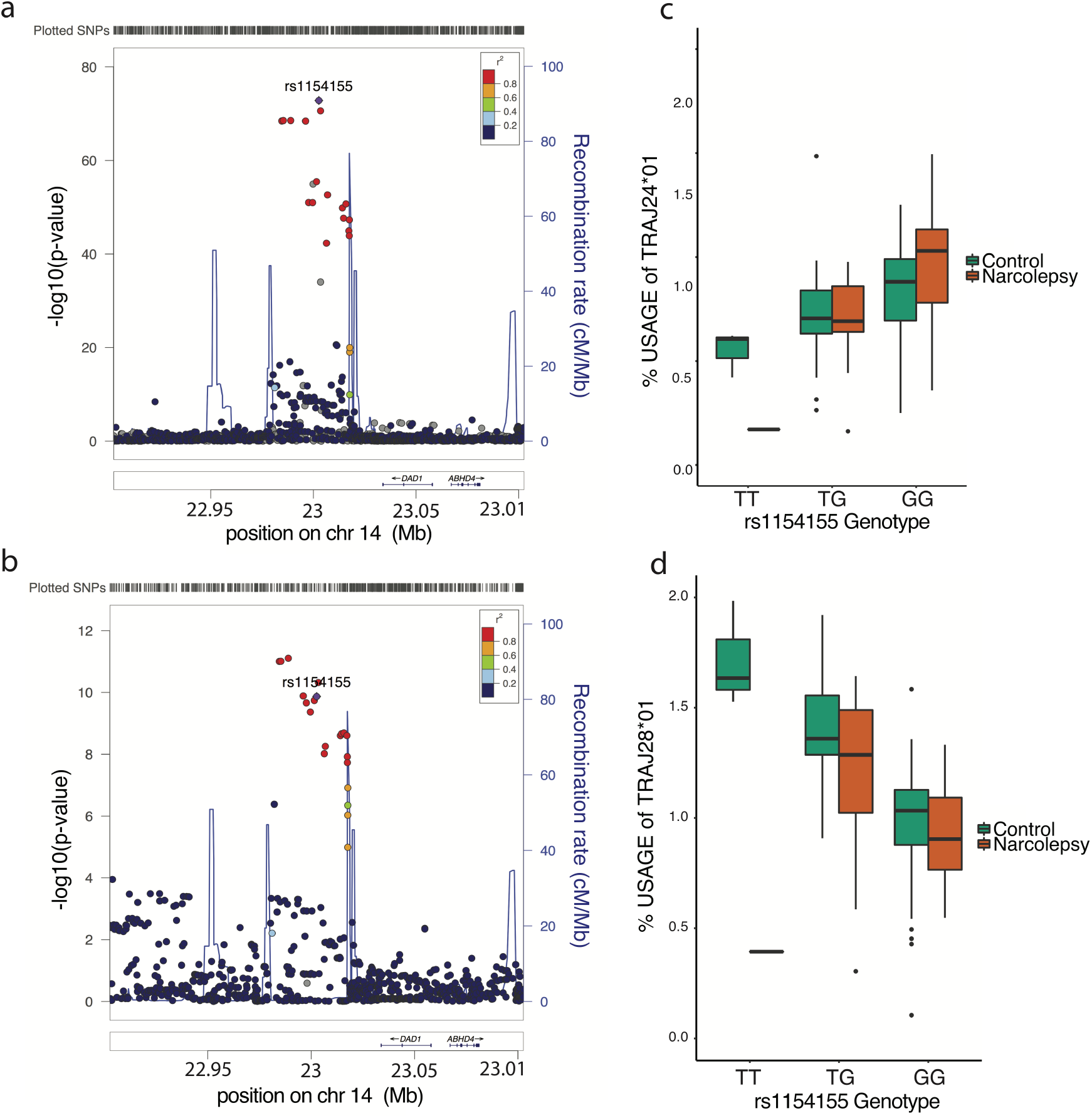
TRA lead variant rs1154155 is associated with repertoire usage of TRAJ24 and TRAJ28 genes. (**a**) T1N association with TRA. T1N association with T cell receptor alpha chain locus spans a region that contains 5 SNPs with almost perfect LD (rs1154155, rs1483979, rs3764159, rs3764160) and high LD over 18kb. (**b**) Usage of TRAJ28*01 in 895 individuals shows similar association with T1N lead variant rs1154155 with posterior probability of 0.958 between narcolepsy and TRAJ28 usage. T cell receptor sequencing in CD4+ T memory cells in 60 type-1 narcolepsy patients and matched controls confirmed the effect of rs1154155 on usage of both (**c**) TRAJ24*01 and (**d**) TRAJ28*01 with higher effect seen in the type-1 narcolepsy cases.

To further investigate the mechanism of the TRA variants specifically on CD4^+^ T cells, which are the most likely causal cell type because of their interactions with DQB1*0602, we performed T cell receptor sequencing of CD4^+^ memory T cells in 40 individuals with T1N and 61 DQ0602 matched controls (**Fig. 3**). Although we found no significantly over-represented T cell clones, we discovered a similar effect of rs1154155 on J28 usage in CD4+ in T1N and healthy controls (beta = −0.32, p-value<0.001, **Fig. 3**). Furthermore, the effect was stronger in individuals with T1N that had significantly lower expression level of TRA J28 than healthy controls (beta = - 0.20, p-value = 0.027). Similarly, the effect of rs1154155 on J24 usage was also similar population cohort (beta = 0.33, p-value<0.001). We also confirmed that these effects were *cis* mediated, and the ratio of J24*01 (F) over J24:02(L) was only 0.4 in heterozygotes, indicating lower allele specific expression with F-narcolepsy associated alleles, with similar effects in other T cell subpopulations (**Supplementary Fig. 10**). The findings suggest that the predisposition to T1N is mediated either by decreasing usage of TRA J28, or by increasing TCR recognition through J24*01, although in this case the effect would be mitigated by decreased expression of this allele.

Within the TRB region, rs1108955 was the leading variant for TRBV4-2, TRBV3-1 and TRBV2 expression (Supplementary Table 6). While it has been observed that individual variants can affect multiple target genes 75, the strongest evidence was seen with TRBV4-2. The leading T1N variant was in perfect LD with the lead variant for TRBV4-2 expression, and the association of same variants for eQTLs in TRB expression for TRBV4-2, TRBV3-1 and TRBV2 pp>0.95 with strongest evidence for TRBV4-2 usage pp=0.99 (**Supplementary Fig. 11**).

We finally examined whether usage of specific TRAJ, TRAV, TRBJ or TRBV genes in CD4^+^ T cells was associated with seasonal influenza vaccination (12 cases versus 5 cases) or with narcolepsy case/control status (59 narcolepsy cases versus 47 DQ0602 controls). Unique T cell receptor gene usage was not associated with influenza vaccination (**Appendix 1, Table 1-16**). However, we did see a statistically significant difference between narcolepsy and controls with *TRBJ1-3*01* usage (p=0.0012, beta=0.00425). Similarly, although TRAJ28 was the second most significantly associating clone between narcolepsy and control with both protective and predisposing clones the association was not statistically significant (p<0.0001, corrected p=1, Appendix 1. Table 23 and Table 24). These findings are in line with usage effects seen with narcolepsy risk variants.

To summarize, the finding that specific TRA and TRB variants associate with narcolepsy suggests specificity for the autoimmune pathology through the T cell receptors. The co-localization of signal at the population sample with expression suggests a direct effect on the specific usage of TRAJ28 expression coding effect on TRAJ24 (F8L) variation as well as TRBV4-2 gene expression. This was also is seen specifically in T cell receptor sequencing in CD4+ T cells and is stronger in patients (p<0.05) suggesting for direct causal effect for disease pathophysiology through expression and autoantigen recognition.

### Multi-loci association of narcolepsy within the HLA region

The strongest association in narcolepsy is within the HLA locus. Strikingly, T1N is one of the few diseases where nearly all affected individuals carry at least one copy of exactly the same HLA allele, DQB1*06:02^5,6^. To fine map this association, we imputed HLA haplotypes using HIBAG 76 and HLA IMP:02^77^. We then performed ethnic specific HLA association and combined them using fixed effects meta-analysis. As expected^5,6^, the strongest association was with the *DQA1*01:02~DQB1*06:02* (DQ0602) haplotype.

To look for additional independent signal, we performed conditional analysis using stepwise forward regression. We detected (1) a strong protective effect of *DQA1*01:01* and *DQA1*01:03* alleles (OR=0.30, p-value<10^−15^ and OR =0.30, p-value<10^−20^, respectively) with combined protective OR=0.41, p-value<10^−40^; (2) predisposing effects for *DQB1*03:01* and *DQA1*01:02* across ethnic groups as shown before ^5,6,78,79^(OR=1.36, p-value<5*10^−8^ and OR=1.68 p-value<5*10^−8^, respectively) (**Supplementary table 7**). The protective effects of DQA1*01:01 and DQA1*01:03 have been suggested to be mediated via heterodimerization with DQB1*06:02, indirectly reducing *cis* encoded DQA1*01:02/DQB1*06:02 (DQ0602) heterodimer availability^5,79^.

Controlling for both *DQB1* and *DQA1* effects, a strong protective association was seen with *DPB1*04:02* allele (p-value<10^−20^) whereas smaller predisposing effect was found with *DPB1*05:01* allele, a mostly Asian subtype (p-value <10^−3^). Finally, after adjusting for the DQ and DP effects significant associations were seen at HLA class I with *A*11:01*, *B*51:01*, *B*35:01* and *B*35:03* and with *A*03:01* (p-value <0.01, Supplementary table 7). These findings confirm and extend results of two previously publications^6,81^, with effects of *B*51:01* likely secondary to LD with A*11:01 in whites.

## Discussion

In this study, we explored genetic risk for narcolepsy and potential disease mechanisms of identified genetic risk factors. The strongest associations were seen with the HLA region. In addition, we confirmed six previously described risk loci (*TRA*, *TRB*, *CTSH*, *IFNAR1*, *ZNF365* and *P2YR11)* and discovered five novel associations in *PRF1*, *CD207*, *SIRPG*, *IL27* and *ZFAND2A.* Analysis of functional consequences of these loci in a multi-ethnic sample discovered remarkable association with immune loci evidenced by individual associations and partitioned heritability enrichment. A notable example is the effect of both missense and regulatory variants in the TRA and TRB regions that had a substantial effect on the T cell receptor chain usage. All these findings strongly suggest specific risk factors in genes controlling immune reactions.

Two loci in addition to the HLA region were implicated in vaccination-associated narcolepsy (*TRA, SIRPB1*). Findings indicate that although genetic factors predisposing to regular and vaccine-triggered narcolepsy are largely shared, there are slight differences. These findings may reflect a primary role for genetic factors in immune response per se versus infection and immune response in other cases. A detailed analysis of the loci where the leading variants for T1N are located suggests both antigen presentation and recognition. Indeed, the majority of variants have effects in antigen presenting cells (*HLA, CTSH*), e.g. dendritic cells (*IFNAR1*, *CD207)*, T cells (*TRA*, *TRB*, *P2YR11, SIRPG),* e.g. T helper cells (*HLA-DQ, HLA-DP, IL27)*, and cytotoxic T cells (*HLA-A*, *PRF1*), sketching a remarkably narrow disease pathway (**Fig. 4**). Accordingly, a direct effect of TRA and TRB associations with T cell receptor expression was seen; TRA lead variant was an eQTL for TRAJ28 and TRAJ24 expression whereas strongest eQTL effect for TRB lead variant was seen with TRBV4-2. The effect was accentuated in T1N cases, suggesting for the first time that specific T cell receptor chains such as TRAJ24, TRAJ28 and TRBV4-2 are strong risk factors for narcolepsy and potentially causal factors recognizing and binding the autoantigen. This association is unique to T1N and has not to our knowledge been seen with other autoimmune diseases.

**Fig. 4.**
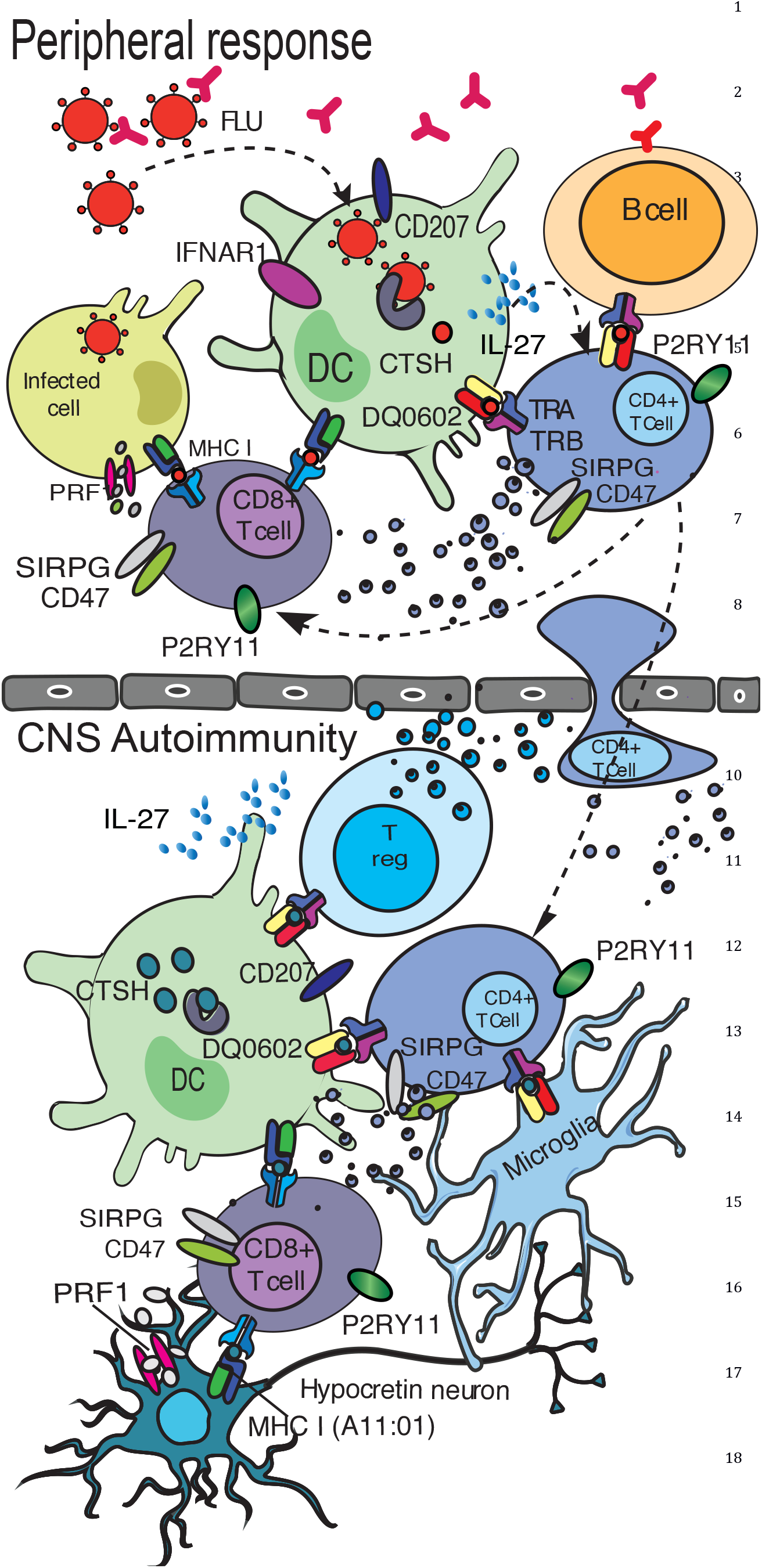
Postulated disease mechanisms in autoimmune narcolepsy. <u>1) Peripheral response:</u> Influenza virions or vaccine protein debris are ingested by DCs facilitated by CD207; flu proteins are processed by cathepsins CTSH and CTSC for presentation by HLA molecules to specific TCRα/β bearing CD4+cells, initiating an immunological synapse and responses to influenza. Presentation by DC is modulated by IFNAR1 in the context of influenza infection. Cross presentation of influenza antigens processed via the MHC class I pathway in DCs is necessary activate CD8+ cells that mature into cytotoxic lymphocytes (CTLs), initiating cell killing of viron infected cells. Activated CD4+ cells produce cytokines such as IFNγ, IL-2 and IL27 which augment cytotoxic activity of CTLs via perforin (PRF1). On the other hand, activated CD4+ cells interact with B-cells via the MHC class II pathway and initiate influenza-specific antibody production, class switching and somatic hypermutation. SIRPG and P2RY11 on activated T cells may also promote cell-cell adhesion and proliferation in this response. <u>2) CNS Autoimmunity:</u> Activated and primed specific CD4+ cells migrate to the CNS where they interact with microglia and resident DCs via DQ0602 bound to an influenza-mimic autoimmune-epitope (derived from hypocretin cells) initiating a secondary memory response. Hypocretin cell proteins are processed by cathepsins CTSH and CTSC for presentation by DQ0602 to specific TCRα*/*β bearing CD4+cells, initiating an immunological synapse and autoimmune responses. Chain usage for TRAJ24-2, TRAJ28, and TRBV4-2 is associated with narcolepsy risk and may be crucial for autoantigen recognition. Further, cross presentation by resident DCs and microglial cells activate specific CD8+cells via MHC class I binding of another hcrt neuron-derived peptides. These primed cytotoxic CD8+ then kill hcrt neurons after recognizing MHC class I (such as A*11:01, associated with narcolepsy independently of DQ0602) bound cognate hcrt neuron derived peptide on hcrt neurons. SIRPB1 on DC or microglia and SIRPG plus P2RY11 on activated T cells may also promote cell-cell adhesion and proliferation in this response. The role of ZFN365 and ZFAND2A is unknown.

In addition, a strong functional connection with Influenza A infection in dendritic cells was found at *IFNAR1,* furthering the role of this virus as a common trigger for the disease. We also discovered associations with *ZNF365* and *ZFAND2A,* ubiquitously expressed transcription factors with, in the case of *ZNF365*, strong known associations with other autoimmune diseases^82,83^. The *ZFAND2A* association (also called Arsenite-inducible RNA-associated protein AIRAP) is unique to narcolepsy, and was opposite in post vaccination cases, an effect that could suggest differential effects on influenza infection and immune response modulation. The *ZFAND2A* associated SNP, is in perfect linkage disequilibrium (r^2^=1) with a very large number of SNPs over a 250 kb region that encompasses and regulates many genes. Of possible interest in this region is *GPR146*, a gene highly enriched in unstimulated macrophages and dendritic cells, whose reduced expression is associated with the INF_γ_ response and suppresses HCMV replication in infected dendritic cells^84^. We were able to examine for the effects of these variants in post Pandemrix^®^ cases. *TRA* association was particularly strong, suggesting involvement of T cell receptor oligoclonallity in autoantigen recognition.

Based on these observations, we propose that narcolepsy is the result of an autoimmune process triggered primarily by influenza-A on an HLA-DQA1*01:02~DQB1*06:02 (DQ0602) background. The involvement of influenza-A is likely to explain why the genetic associations we found are universal. Indeed, influenza is one of few viruses that act worldwide on a seasonal basis. The universal association is especially clear for DQ0602 as it is found with different HLA-DRB1 alleles, DRB1*15:01 in White (Europe and USA) and Asians (China, Korea, Japan and India), but DRB1*15:03 or DRB1*11:01 in Blacks (confusion of ancestral continent of origin and sample location?) ^5,6^. The primacy of DQ0602 over DRB1*15:01 is also demonstrated by the fact DRB1*15:01~ DQA1*01:03~DQB1*06:01 haplotype is not associated with narcolepsy in China and by the fact additional DQ effects are mostly mediated by DQA1 alleles that interact in trans with DQB1*06:02. In contrast to narcolepsy, other autoimmune diseases commonly have different HLA associations or disease presentations across countries, and resulting HLA associations are more complex. Type 1 diabetes, for example, is well known to be primarily associated with HLA-DQ in Whites whereas DRB1*04:05 specific effects are evident in Japan where the disease is rare^83,85,86^ ^77^.

Other autoimmune diseases, unlike narcolepsy, are also associated with a plethora of autoantibodies and known autoantigen targets. For example Insulin, GAD, IA-2 and ZNT8 are involved in T1D and β-cell antigen targeting, suggest that these other diseases involve multiple B and T cell mechanisms and antigens, likely explaining the weaker and more complex HLA effects and a lack of association with any specific TCR polymorphisms. It is our hypothesis that the strong effects of TCR polymorphisms in narcolepsy likely represent the fact autoimmunity in this disease is oligoclonal and limited to one or a few hypocretin cell antigen epitopes. These epitopes may bind DQ0602 specifically and involve a few αβTCR receptors containing TRAJ24, TRAJ28 or TRBV4-2 (**Fig 4**). Other groups have suggested involvement of TRIB2, prostaglandins and HCRTR2^87-91^. However, these associations have not been universal. Systematic studies of T-cell reactivity with TCR identification in the context of DQ0602 and flu or autoantigen epitopes are ongoing in various laboratories to address this issue.

In this study, perforin, a gene of critical importance to NK and CD8^+^ T cell cytotoxicity was strongly protective of narcolepsy, whether or not it was triggered by vaccination. In the context of compound null heterozygotes of the perforin gene, A91V has been is associated with late onset hemophagocytic lymphohistiocytosis (HLH) type 2^49^, a recessive disorder associated *PRF1* null alleles. HLH type 2 is characterized by excessive T cell activation that may involve abnormal reactivity to viral pathogens ^50^or decreased CD8+ T cytotoxic pruning of dendritic cells^51^. Interestingly, Prf1 knock-out mice do not develop the syndrome unless infected with viruses such as murine lymphocytic chorio-meningitis virus or murine cytomegalovirus, a phenomenon involving CD8^+^ T cells and increased IFNγ^50^. Other perforin-damaging mutations have also been anecdotally associated with susceptibility to multiple sclerosis^52^ and T1D^53^. Importantly, the allele associated with narcolepsy impairs cytotoxicity and cell killing, suggesting that the effect of the variant on cytotoxicity may be targeting hypocretin cells directly.

Although it is conceivable NK cells could be involved, the most likely explanation is involvement of CD8^+^ T cell in hypocretin cell killing in collaboration with CD4^+^ T cells or microglia. This was also supported by CTSC association, an enzyme of critical importance to cytotoxic CD8^+^ activation of pro-granzymes^58^. Bernard-Valnet et al.^92^ used transgenic mice with expression of a neoantigen in hypocretin neurons, and found that infusion of CD8^+^ T cell targeting the neoantigen were able to cause hypocretin cell destruction while infusion of neoantigen-specific CD4^+^ T cell alone was insufficient, although CD4^+^ T cells migrated closely to the target neurons. These earlier experiments together with genetic association with PRF1 variants suggest a direct role of CD8+ T cells in hypocretin cell destruction. CD8+ mediation of cell killing has also been suggested by observation of a CD8 T cell infiltrate in a paraneoplastic anti-Ma2 encephalitis case with symptomatic hypocretin cell destruction ^93^.

In summary, although the culprit autoantigen has not been identified, genetic data indicate autoimmunity in T1N with strongest genetic overlap with T1D, another organ-specific autoimmune disease suggesting shared pathophysiology. A particularity of the disease is involvement of polymorphisms such as in IFNAR1 that regulate response to influenza-A infection, a result that complement epidemiological studies indicating seasonality of disease onset^42^ and increased incidence that has occurred following vaccination with Pandemrix^®^ in Europe ^13,18,19^. Other genetic factors implicate dendritic processing of antigens, presentation by DQ0602 to CD4^+^ T cells and subsequent cell killing of hypocretin neurons by CD8^+^ cells, with likely involvement of only a few autoantigen epitopes and a restricted number of T-cell receptors. The lack of detectable autoantibodies has made objective demonstration of autoimmunity challenging, but will likely made the eventual discovery of the culprit T cell antigen even more informative to our understanding of T cell immunity in the brain.

## Methods

### Study subjects

5,339 unrelated individuals with type 1 narcolepsy^8,9^, and 20,518 ethnicity-matched controls were included in the study. In addition, 245 individuals with vaccination related narcolepsy and 18862 controls were recruited in Finland (N=76 cases and 2796 controls), Sweden (N=39 and 4894 controls), Norway (N=82 cases and 429 controls), and United Kingdom and Ireland (N=48 cases and 10743 controls) ^13,16,94,95^. All cases had documented immunization with Pandemrix^®^. All cases had narcolepsy with clear-cut cataplexy and were *DQB1*06:02* positive, or had narcolepsy with documented low hypocretin-1 in the cerebrospinal fluid. Informed consent in accordance with governing institutions was obtained from all subjects. The research protocol was approved by IRB Panels on Medical Human Subjects at Stanford University, and by respective IRB panels in each country providing samples for the study.

### Genotyping

Subjects were genotyped using Affymetrix Affy 5.0, Affy 6.0^8^, Affymetrix Axiom CHB1^9^, Affymetrix Axiom EUR, Axiom EAS, Axiom LAT, Axiom AFR, Axiom PMRA and Human Core Exome chip platforms. Genotypes were called with Affypipe^96^, Affymetrix genotyping console or Genome Studio. Markers with genotyping quality (call rate < 0.95) or deviation from Hardy-Weinberg equilibrium (p-value<10^−6^) were discarded from further analysis. Samples were checked for relatedness with filtering based on proportion of identity-by-descent using cut off >0.2 in PLINK 1.9 PI_HAT score ^88^. One pair of related individuals was removed. If related individuals were a case and a control, cases were retained in the analysis. Three first principal components within each cohort were visualized and outliers were removed. **Supplementary Table 1** shows for each cohort N QCed original genotypes, N for those passing the QC and N for individuals removed during QC.

### Imputation

We imputed samples by prephasing cases and controls together using SHAPEIT v2.2^89^ and imputed with IMPUTE2 v2.3.2 ^97,98^ and 1000 genomes phase 1v3 build37 (hg19) in 5Mb chunks across autosomes. For variants having both imputed and genotyped values, the genotyped values were kept except for those individuals where the genotype was missing. In this case imputed values were kept.

### Analysis

Analyses for all data sets were performed at Stanford University except for the Finnish and Swedish vaccination related cases and European Narcolepsy Network samples, which were analyzed by respective study teams using exactly the same analysis. Genome-wide association analysis was first performed in each case control group separately using SNPTEST v.2.5.2^99^. We used linear regression implemented in SNPTEST method score adjusting for ten first principal components in order to adjust for cohort specific population stratification. Standard post imputation quality control was done: Variants with info score <0.7 and minor allele frequency (MAF) <0.01 were removed from the analysis. Signals specific for one genotyping platform only and variants in each locus with heterogeneity p-value<10^−20^ were removed. We used fixed effects model implemented in METAv1.7 with inverse-variance method based on a fixed-effects model for combining the association results^100^. In total 12,600,187 markers across the studies were included in the final case control meta-analysis. Significance level for statistically significant association was set to genome-wide significance (p-value<5*10^−8^) controlling for multiple testing. Overall test statistics showed no genomic inflation. GCTA was used for heritability and gene based tests ^101^. Coloc analysis was done using coloc package in R version 3.4.2 (2017-09-28) ^40^, Manhattan and QQ-plots were created with QQman or FUMA ^97^. Shared heritability was estimated using LD score regression^32^.

### Typing and imputation of HLA variants

High resolution HLA imputation in 4-digit resolution (2-field, amino acid level) for HLA *A, B, C, DRB1, DQA1, DQB1*, *DPA1* and *DPB1* was performed using HLA*IMP:02 as implemented in Affymetrix HLA or the HIBAG package in R version 3.1.2 (2014-10-31). HIBAG is an HLA imputation tool that uses attribute bootstrap aggregation of several classifiers (SNPs) to select groups of SNPs that predict HLA type and allows the use of own HLA reference panels^76^. Reference HLA types were used from published imputation models and for Asian and Blacks obtained with Sirona sequencing ^102^in ethnic specific populations N=500 Blacks, N=2,000 Whites and N=368 Asians. Imputation accuracy was further verified by Luminex HLA typing in a subset of samples and accuracy was over 95% for all ethnic groups and common alleles with > 5% frequency in population. For all alleles the accuracies were for Whites: 0.98 in HLA-A, 0.97 in HLA-B, 0.98 in HLA-C, 0.96 in HLA-DRB1, 1.00 in HLA-DQA1, 1.00 in HLA-DQB1, 1.00 in HLA-DPA1, and 0.92 in HLA-DPB1 and for Asian for alleles where typing was also available 0.95 for HLA-DRB1, 0.94 for HLA-DQA1, and 0.98 for HLA-DQB1.

### Analysis of HLA variants

HLA effects in narcolepsy were analyzed as described before^6^. We examined altogether variation from 23,410 individuals with 9,789 Asians, 13,621 Whites. In each ethnicity HLA alleles were analyzed using additive model under logistic regression adjusting for 10 first population specific principal components to adjust for local population stratification. We identify independent associations using conditional analysis (stepwise forward regression in each cohort). Fixed effects meta-analysis was used to combine associations using Plink 1.9^103^ and R version 3.2.2. We considered alleles sustaining Bonferroni correction for correction of number of alleles with minor allele frequency over 2% (N=110 HLA alleles) significant resulting in Bonferroni cut-off p=0.00045.

### Analysis of expression quantitative trait loci (eQTL)

We used tissue specific summary statistics from the GTEx consortium and from Westra et al. to examine total blood specific effects of associating variants on gene expression^75,104^. Furthermore, we examined how the genetic variants modulated T cell and antigen presenting (dendritic cell and monocyte) gene expression by RNA sequencing and RNA expression. To examine environment specific triggers for eQTLs we challenged the dendritic cells on influenza-A infection, or stimulated them with interferon or LPS^105,106^. Finally, we identify short range (cis) SNPs and trans HLA alleles association with TCR V and J usage estimated from total peripheral blood RNA sequencing as described before^73^.

### T cell receptor RNA sequencing in matched narcolepsy case control data set and in population cohorts

We performed RNA sequencing in 895 individuals with total blood RNA sequencing and in T cells from 60 individuals with narcolepsy and 60 healthy individuals from using total CD4+ T cells, CD4+ T memory and CD8+ T cell populations. We used fastqc to infer quality and trimmed low quality reads. We then performed barcode demultiplexing, after which local blast was used to align and extract CDR3s. Linear regression was fit for TRA usage ~ Genotype adjusting for age and gender, RNA sequencing lane and case/control status as covariates. We also analyzed separately coding consequences for each TRAJ24 containing productive CDR3 fragment as one of the most significantly associating SNPs was a coding SNP (rs1483979) was changing an amino acid Leucine to Phenylalanine. These ‘LQF’ and ‘FQF’ were extracted and their frequencies were computed. Ratio of FQF/(LQF+FQF) was further computed across all the samples.

## Acknowledgements

We want to thank Dr. Mehdi Tafti for identifying overlapping samples from EU-NN and those genotyped at Stanford University. We want to thank GlaxoSmithKline (GSK) for the collaboration providing funds to examine the effect of seasonal influenza vaccination and T cell receptor sequencing in narcolepsy (Appendix 1). We thank Briana Gomez for her valuable contribution in editing the manuscript. This study was primarily supported by grants from wake up narcolepsy, Jazz Pharmaceutical, donations from narcoleptic patients, and a previously funded NIH-23724 grant (EM). It was also supported by Academy of Finland (#309643), Sigrid Juselius Foundation, Finnish Cultural Foundation, Orion Farmos Research Foundation, Instrumentarium Science Foundation, Jalmari and Rauha Ahokas Foundation (HMO). This work was also finally supported by the United States Department of Defense (DoD) through the National Defense Science & Engineering Graduate Fellowship (NDSEG) program and by Stanford University through the Stanford Graduate Fellowship program. UK Biobank Resource has been used for this study. We would like to thank the participants and researchers from the UK Biobank who contributed or collected data. The work has been supported by National Institute of Mental Health Grant 5RC2MH089916 for the Depression Genes and Networks Project. The Swedish post-pandemrix Narcolepsy genetics, with Tomas Olsson and Ingrid Kockum as PIs, received grant support from the Swedish medical product agency. Han Fang was supported by Ministry of Science and Technology (2015CB856405) and NSFC (81420108002,81670087).

